# Gallionellaceae in rice root plaque: metabolic roles in iron oxidation, nutrient cycling, and plant interactions

**DOI:** 10.1101/2023.04.07.536048

**Authors:** Clara S. Chan, Gretchen E. Dykes, Rene L. Hoover, Matt A. Limmer, Angelia L. Seyfferth

## Abstract

On the roots of wetland plants such as rice, Fe(II) oxidation forms Fe(III) oxyhydroxide-rich plaques that modulate plant nutrient and metal uptake. The microbial roles in catalyzing this oxidation have been debated and it is unclear if these iron-oxidizers mediate other important biogeochemical and plant interactions. To investigate this, we studied the microbial communities, metagenomes, and geochemistry of iron plaque on field-grown rice, plus the surrounding rhizosphere and bulk soil. Plaque iron content (per mass root) increased over the growing season, showing continuous deposition. Analysis of 16S rRNA genes showed abundant Fe(II)-oxidizing and Fe(III)-reducing bacteria (FeOB and FeRB) in plaque, rhizosphere, and bulk soil. FeOB were enriched in relative abundance in plaque, suggesting FeOB affinity for the root surface. Gallionellaceae FeOB *Sideroxydans* were enriched during vegetative and early reproductive rice growth stages, while a *Gallionella* was enriched during reproduction through grain maturity, suggesting distinct FeOB niches over the rice life cycle. FeRB *Anaeromyxobacter* and *Geobacter* increased in plaque later, during reproduction and grain ripening, corresponding to increased plaque iron. Metagenome-assembled genomes revealed that Gallionellaceae may grow mixotrophically using both Fe(II) and organics. The *Sideroxydans* are facultative, able to use non-Fe substrates, which may allow colonization of rice roots early in the season. FeOB genomes suggest adaptations for interacting with plants, including colonization, plant immunity defense, utilization of plant organics, and nitrogen fixation. Together, our results strongly suggest that rhizoplane and rhizosphere FeOB can specifically associate with rice roots, catalyzing iron plaque formation, with the potential to contribute to plant growth.

**Importance:** In waterlogged soils, iron plaque forms a reactive barrier between the root and soil, collecting phosphate and metals such as arsenic and cadmium. It is well established that iron-reducing bacteria solubilize iron, releasing these associated elements. In contrast, microbial roles in plaque formation have not been clear. Here we show that there is a substantial population of iron oxidizers in plaque, and furthermore, that these organisms (*Sideroxydans* and *Gallionella*) are distinguished by genes for plant colonization and nutrient fixation. Our results suggest that iron-oxidizing and iron-reducing bacteria form and remodel iron plaque, making it a dynamic system that represents both a temporary sink for elements (P, As, Cd, C, etc.) as well as a source. In contrast to abiotic iron oxidation, microbial iron oxidation results in coupled Fe-C-N cycling, as well as microbe-microbe and microbe-plant ecological interactions that need to be considered in soil biogeochemistry, ecosystem dynamics, and crop management.

## Introduction

Iron cycling is a key biogeochemical process in rice paddies that plays important roles in the growth and quality of rice crops. Fe(II) oxidation leads to the production of Fe(III) oxyhydroxides (ferrihydrite, goethite, lepidocrocite), which are strong sorbents of organic carbon, phosphate, and metal(oid)s such as As and Cd (1–5); Fe(III) reduction dissolves the oxyhydroxides and releases sorbed nutrients and toxins (6). Of particular interest are iron oxyhydroxide coatings (plaques) that develop on the surface of rice roots (rhizoplane), as well as in the rhizosphere (satellite plaque). Fe(II)-oxidizing and Fe(III)-reducing bacteria (FeOB and FeRB) have been documented in association with a variety of wetland plants (7, 8). Because plaque is closely associated with rice roots, its formation and dissolution may limit or drive plant uptake of oxyhydroxide-sorbed chemicals (9–12), motivating us to study mechanisms of rice root plaque formation. FeOB can catalyze formation of Fe(III) oxyhydroxides via their metabolism. FeOB have been detected in paddy soil (11, 13, 14), making them a potential mechanism for iron plaque formation in conditions in which they can outcompete abiotic Fe(II) oxidation.

The rice rhizoplane is in fact an ideal niche for microaerophilic FeOB, which gain energy by coupling Fe(II) oxidation to oxygen reduction, and thus grow best where Fe(II) and O_2_ fluxes are high. Although Fe(II) oxidation coupled to nitrate reduction is possible in theory, it is unlikely to be a significant process because N is a limiting nutrient throughout most of the rice growing season (15–18). Rice paddy soil is rich in Fe(II) formed by microbial Fe(III) reduction, thus plaque formation is partially dependent on activities of FeRB. Rice root aerenchyma are conduits of O_2_, which diffuses from roots into saturated soil rich in Fe(II), providing both electron donor and acceptor for FeOB. Aerobic FeOB must compete with abiotic oxidation, and kinetics studies showed they become the dominant mechanism of iron oxidation as oxygen concentrations decrease (19, 20). Different neutrophilic FeOB can thrive at O_2_ concentrations <1-100 μM (21–24), concentrations that coincide with ranges typically found at the surface of rice roots (few micromolar to tens of micromolar O_2_) (25, 26). Thus, it is plausible that a significant proportion of plaque iron oxidation is microbial.

Indeed, studies have increasingly documented FeOB in rice paddy soil, thus increasing recognition that microbes can contribute to iron oxidation in this environment. FeOB can be documented by culturing, 16S rRNA gene analyses, and metagenomic studies, which give different evidence of potential contributions. Numerous studies have shown that FeOB can be cultured from rice paddy soil, with a range of taxa represented (13, 24, 27–31). While culturing demonstrates microbial iron oxidation activity, it only proves that soil-derived organisms are capable of iron oxidation, but cannot show that these FeOB represent the major active organisms *in situ*. Studies using 16S rRNA genes give a broader view of soil microbial communities, but it is often unclear which organisms are FeOB, or identification is equivocal due to the metabolic flexibility of putative FeOB. The exceptions are taxa specifically known for iron oxidation, like Gallionellaceae iron-oxidizing genera, including *Gallionella*, *Ferrigenium*, and *Sideroxydans* (28, 32, 33). As their primary metabolism is iron oxidation, detection of Gallionellaceae FeOB in field samples is a strong indication of microbial iron oxidation in the environment.

Gallionellaceae FeOB have been cultured from paddy soil, including the isolates *Gallionella/Ferrigenium kumadai* An22 (28, 34) and *Sideroxydans* sp. (24). While the isolate An22 was named *Ferrigenium* based on 16S rRNA gene dissimilarity from *Gallionella*, subsequent full-genome analyses showed that it is a *Gallionella* (35). Gallionellaceae have also been identified using 16S rRNA gene analyses including universal and taxa-specific primers in paddy soil and the rice rhizosphere (31, 36–39). Their abundance has been shown to correspond to iron oxidation in paddy soil and soil incubations (30, 40). *Gallionella* have been detected in rhizoplane samples by qRTPCR (41). However, it has not been shown whether Gallionellaceae are specifically enriched in abundance at the rhizoplane (vs. soil) and associated with plaque oxyhydroxides. Schmidt and Eickhorst (2014) used CARD-FISH to show Betaproteobacteria (which includes Gallionellaceae) on root surfaces, and suggested that these may represent Gallionellaceae detected via 16S rRNA sequences (42). In all, there is strong evidence for the existence of these FeOB in paddy soil, but it is still unclear if Gallionellaceae are specifically associated with plaque and therefore contribute to iron oxyhydroxide formation. If FeOB are in fact associated with plaque, this close association with the root could cause them to compete with the plant for nutrients like N, or alternatively, FeOB may provide fixed N and other nutrients to help promote plant growth. To understand the biogeochemical cycling in the rice root rhizosphere, we need to determine the dynamics and functions of plaque microorganisms, and their interactions with the rice plant.

In this study, we investigated the spatial and temporal dynamics of rice-associated FeOB and FeRB in rice paddies over a growing season, using 16S rRNA gene analyses of bulk soil, rhizosphere, and plaque samples. This was coupled to plaque Fe analyses as well as examination of metagenomic-assembled genomes of FeOB to more specifically identify metabolic capabilities and niche. Together, this allowed us to identify the major FeOB enriched in iron plaque and gives insight into how microbes contribute to plaque formation over the life cycle of field-grown rice.

## Results and discussion

### Rice paddy experiment

Rice was grown in the University of Delaware (UD) Rice Information, Communication, and Education (RICE) Facility on the UD Farm in Newark, Delaware. For this study, rice was grown in 12 soil-filled paddy plots (2×2 m), or mesocosms, each containing 49 plants, in either unamended native soil, or soil that had received Si amendments a year before this experiment (Fig. S1). Details on soil chemistry and treatments, including fertilization, and water management can be found in the Methods section. Previous studies of the microbial community showed that Si amendments had less of an impact than redox status (98, 99). Given the variability in chemistry and community composition across the 12 paddies, here we show chemistry (plaque iron content and mineralogy) for each paddy individually (43), and then aggregate samples from all treatments to evaluate the most abundant FeOB and FeRB across all conditions.

### Plaque Fe content and mineralogy

We measured plaque Fe content on roots at five timepoints over the growing season (See Methods and Figure S1 for details of rice paddy experimental setup). Plaque Fe was quantified as dithionite-citrate-bicarbonate (DCB)-extractable Fe per dry root mass, obtained by removing one whole plant per paddy at each time point. The amount of plaque Fe per dry root mass increased over time, with the highest rate of plaque increase between heading to grain maturity (71-88 days past transplant (DPT) and 88-98 days DPT, **Fig. 1a**). The increase in plaque Fe is not a function of increasing Fe in porewater, as dissolved Fe(II) did not increase over time (measured in bulk soil; **Fig. 1b**). The continuous increase of plaque Fe over time shows active formation of plaque throughout the growing season.

**Figure 1.**
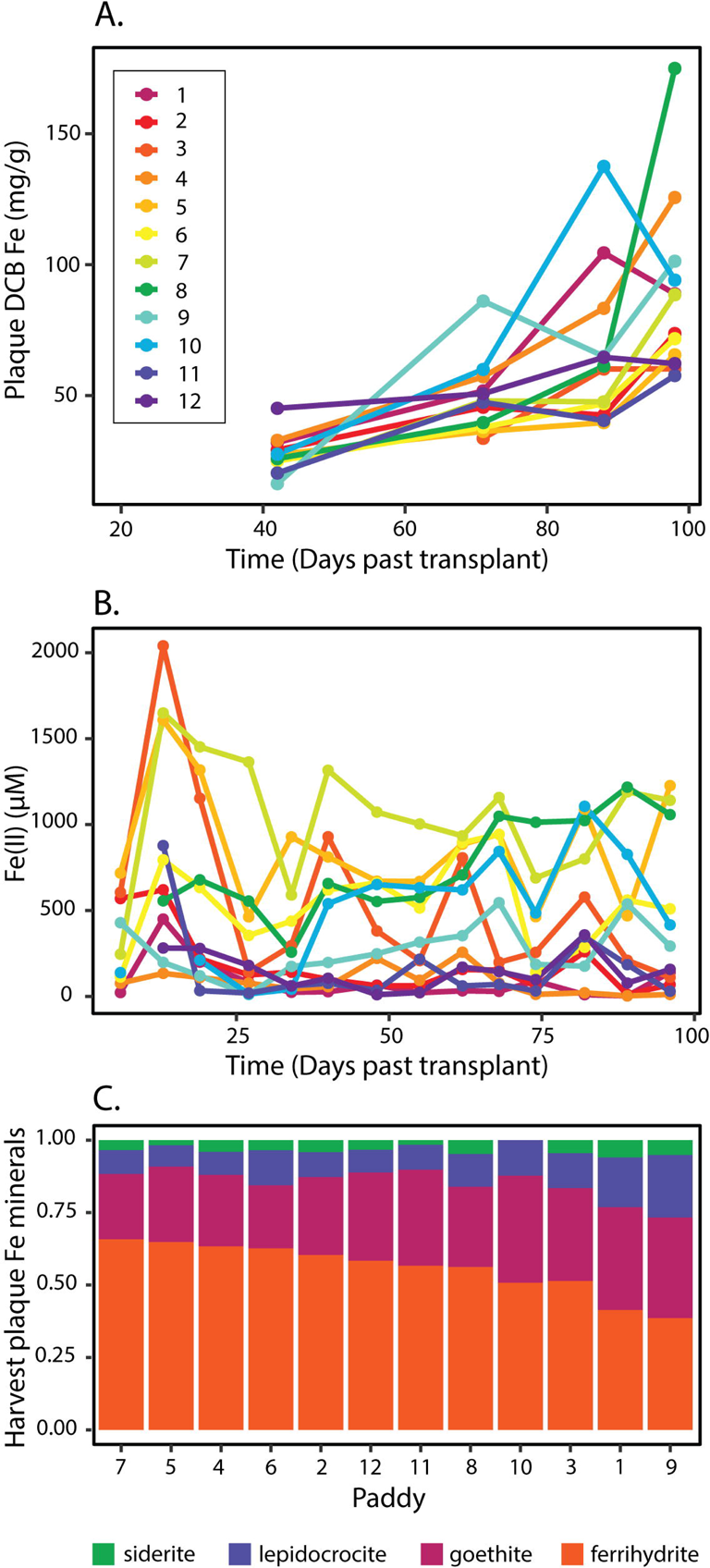
Plaque iron content compared to porewater Fe(II), and mineral composition. (A) Plaque iron content per dry root mass over the growing season plotted for 12 paddies sampled. (B) Porewater Fe(II) over the growing season plotted by and colored by paddy (see legend in A). (C) Plaque Fe mineral composition at harvest, by paddy, in order of decreasing ferrihydrite.

On initial precipitation, Fe(III) oxyhydroxide minerals tend to be poorly crystalline phases like ferrihydrite, often increasing in crystallinity as they age. Thus, to evaluate the potential for active plaque formation, we assessed mineral composition of iron plaque at harvest (98 DPT; dataset from (43)). The iron plaque mineral composition at harvest was primarily ferrihydrite, with the plaque ranging from 38.6%-65.7% ferrihydrite (average 55.9%), followed by goethite (21.8-37.7%, average 29.4%) (**Fig. 1c**). These results are consistent with previous observations that ferrihydrite is the most abundant iron plaque mineral in rice plants at harvest (9, 10, 44–47), and other wetland plants (48, 49) with goethite and lepidocrocite as a substantial proportion (10, 44, 45, 47–50).

While the presence of ferrihydrite in the plaque at harvest is consistent with active plaque formation until harvest, ferrihydrite minerals are sometimes stabilized by organics (e.g. root exudates) or inorganic ions (e.g. silicate), which are present in the soil solution throughout the growing season (9, 51, 52). Here, nine out of twelve paddies were amended with additional silicon in various forms while three control paddies (nos. 1, 9 and 10) had no silicon added. Silicon addition is known to retard the crystallization of ferrihydrite to higher ordered phases (51, 53). Although plaque from the control paddies generally had lower ferrihydrite than the others at harvest, ferrihydrite still comprised at least 38% of the Fe mineral composition in plaque at harvest, even in non-amended controls. When considered together, increasing plaque Fe and ferrihydrite in the plaque at harvest, regardless of treatment, imply active precipitation of Fe oxyhydroxides over the course of the growing season.

### Assessing the Fe-cycling microbial community by 16S rRNA gene analyses

#### FeOB and FeRB in iron plaque

We assessed the community composition using the 16S rRNA gene V4-V5 region. We sampled bulk soil, rhizosphere soil, and Fe plaque from 12 rice paddy plots over the course of one growing season (5 time points). In total, we analyzed 130 samples, from which we retrieved a total of 37,072,594 quality-filtered sequences, with rarefaction analysis showing sufficient sequencing depth (**Fig. S2**). The sequences clustered into 11,678 operational taxonomic units (OTUs, 97% similarity), of which 3,914 were assigned taxonomy to the genus level (**Table S1**), from which we could predict FeOB and FeRB.

Here, we focus on organisms associated with the iron plaque. To determine the most abundant plaque community members, we ranked the OTUs by their median relative abundance in the plaque (**Fig. 2a**) and looked for iron cycling organisms (**Table S2**). Twelve OTUs had median abundances greater than 0.5%, including two FeOB OTUs. The most abundant plaque OTU (OTU7) was initially identified as Gallionellaceae, and a BLAST search showed 100% identity to *Gallionella/Ferrigenium kumadai*, an FeOB that was isolated from rice paddy soil (28); hereafter, OTU7 is referred to as *Gallionella*. The third most abundant plaque OTU (OTU2) has 100% identity to a number of sequences, including *Sideroxydans* clones from riparian wetlands (acc. nos. JQ060114, JQ060109, JQ060108; (54)) and sequences from rice paddy soil (AB657736; (55)). *Sideroxydans* is a well-characterized FeOB within the *Gallionellaceae*, and has previously been cultured from rice paddies (56) and wetland plant roots (8).

**Figure 2.**
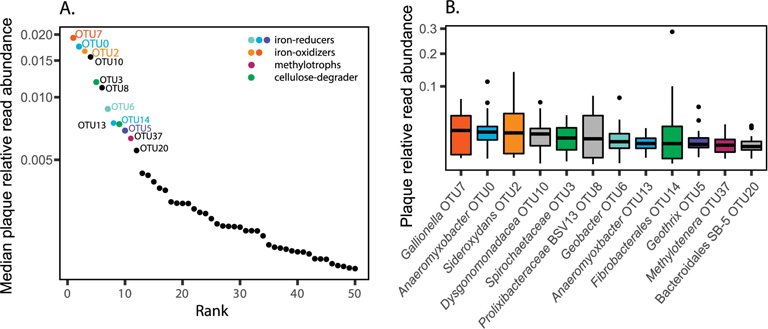
(A) Log 10 median rank relative abundance curve of top 50 plaque organisms. (B) Box and whisker plot showing square-root transformed relative abundance of OTUs with median relative abundance above 0.5% in plaque. Boxes are colored according to known taxonomy with orange/yellow representing FeOB, pink/purple FeRB, blue methylotrophs, and green cellulose degraders. See Tables S1 and S2 for abundance data, ranking calculations, and taxonomic classification of OTUs.

The top OTUs also include several FeRB, including two *Anaeromyxobacter* OTUs, a *Geobacter*, and a *Geothrix* OTU, all of which have been previously identified in paddy soils (57–60). The second most abundant OTU was *Anaeromyxobacter* OTU0, which had 100% identity to an organism in a rice paddy soil enrichment (acc. no. MF547866; (61)). The *Geobacter* and *Geothrix* OTUs were also related to sequences from rice paddy soils (OTU6 100% to MF968163, OTU5 100% to KY287536). In addition, there is an OTU related to *Rhodoferax* (OTU93) that was only classified on the family level, but is 97% similar to *Rhodoferax ferrireducens* T118 and 97% similar to a *Rhodoferax* full length 16S sequence from the 9BH metagenome described below.

In addition to iron-cycling organisms, the highly abundant plaque community members included one methylotroph (*Methylotenera* OTU37) and known cellulose degraders (*Spirochaetaceae* OTU3 and *Fibrobacterales* OTU14, (62–64). Beyond the top 12 OTUs, there were many other iron cycling OTUs, with a notable diversity of FeRB, including 32 *Anaeromyxobacter* and 32 *Geobacter* OTUs detected in plaque. This contrasted with three Gallionellaceae OTUs, suggesting a greater range of iron-reducing niches, despite the comparable abundance of FeOB and FeRB in plaque.

#### Enrichment (increased relative abundance) of iron-cycling microbes in the plaque

If FeOB or FeRB are involved in plaque formation, we would expect to see them specifically enriched in plaque compared to bulk soil, and possibly also enriched in the rhizosphere. The FeOB and FeRB OTUs were generally present in all sampled biospheres: bulk, rhizosphere, and plaque. Thus, we were able to calculate an enrichment factor as the ratio of relative abundance in plaque versus bulk soil or rhizosphere versus bulk soil (**Fig. 3)**. Because cell concentrations are higher in the rhizosphere and plaque relative to bulk soil (42, 65), these represent lower limits on absolute enrichment factors.

**Figure 3.**
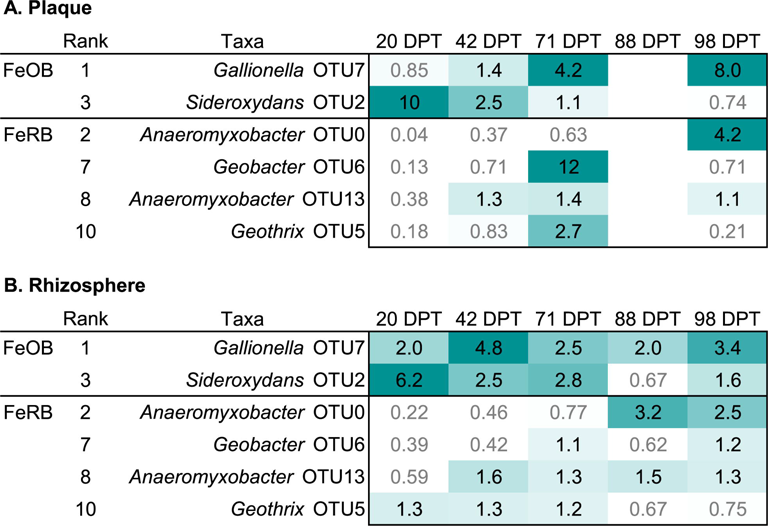
Enrichment factors in plaque (A) and rhizosphere (B) of most abundant FeOB and FeRB OTUs over growing season (DPT=days past transplant). Enrichment factor was calculated as relative abundance in plaque or rhizosphere divided by relative abundance in bulk soil, and the median is given for each timepoint. Boxes are shaded according to enrichment factor, with those below 1 (not enriched) in white with gray text. (No plaque sample available for 88 DPT.) OTUs are ranked by median relative abundance in plaque.

While there was always an FeOB enriched in the plaque, different FeOB OTUs dominated over the growth cycle. The FeOB *Sideroxydans* OTU2 was enriched at earlier time points (**Fig. 3**). *Sideroxydans* OTU2 was most enriched at the first time point (20 DPT), at 10x relative abundance in plaque, relative to bulk soil. *Sideroxydans* OTU2 was still enriched in plaque at early reproduction (42 DPT), but continued to decrease in plaque while increasing in bulk soil (**Fig. 4a**). In contrast, *Gallionella* OTU7 is most abundant and enriched in plaque at grain maturity (98 DPT) at 8x bulk soil abundance, though it was also enriched at early reproduction in the rhizosphere (42 DPT) (**Fig. 3**, **Fig. 4b**). These contrasting enrichment and abundance patterns suggest that *Sideroxydans* and *Gallionella* have different niches, and that *Gallionella* may be more responsive as the plants mature. In all, FeOB are abundant at all time points, and there is always one FeOB enriched in plaque at all times, consistent with iron oxidizer roles in plaque formation.

**Figure 4.**
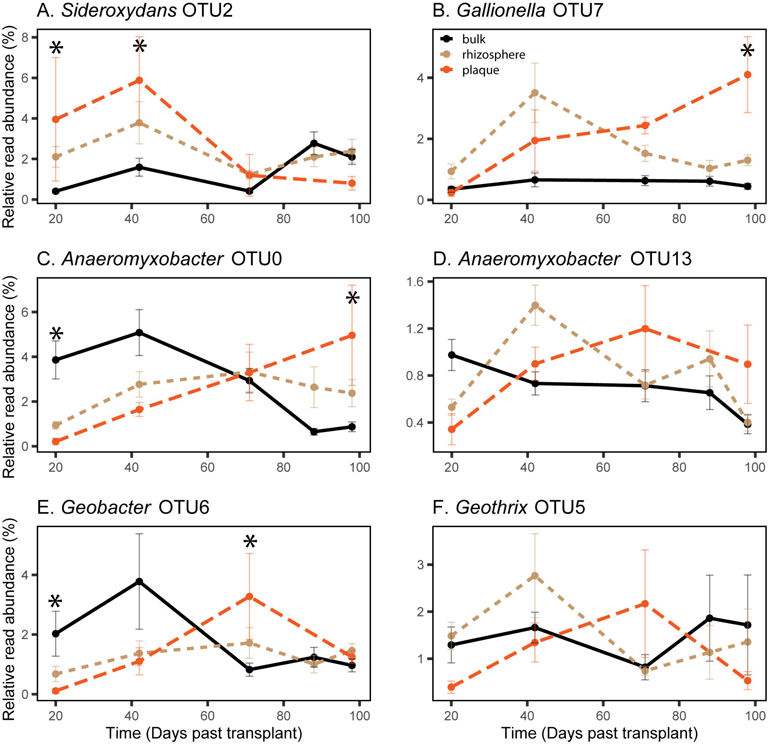
Relative read abundance of top FeOB and FeRB in plaque (with median relative abundance above 0.5%) over time for each biosphere. Abundance in bulk soil plotted with black solid line, rhizosphere plotted in tan short-dashed line, and plaque plotted in orange long-dashed line. Error bars show standard error. Asterisks indicate differential abundance in the plaque and bulk soil where adj. p <0.1 based on DESeq2 analysis. Bulk 20 DPT n=12, rhizosphere 20 DPT n=12, plaque 20 DPT n=3, bulk 42 DPT n=11, rhizosphere 42 DPT n=10, plaque 42 DPT n=6, bulk 71 DPT n=11, rhizosphere 71 DPT n=11, plaque 71 DPT n=4, bulk 88 DPT n=12, rhizosphere 88 DPT n=12, bulk 98 DPT n=11, rhizosphere 98 DPT n=11, plaque 98 DPT n=4.

In contrast to FeOB, the FeRB OTUs are initially low and increase in plaque over time, in both relative abundance (**Fig. 4**) and enrichment factor (**Fig. 3**). Both of the top two FeRB OTUs, *Anaeromyxobacter* OTU0 and *Geobacter* OTU6 are more abundant in the bulk soil at earlier time points (**Fig. 4c, 4e**). *Geobacter* OTU6 later becomes enriched in the plaque (12× bulk) at heading (71 DPT), while *Anaeromyxobacter* OTU0 is most abundant at grain maturity (98 DPT, 4x bulk). *Anaeromyxobacter* OTU13 and *Geothrix* OTU5 are similar to *Geobacter* OTU6 in that they peak in plaque at heading (71 DPT) (**Fig. 3**, **Fig. 4d, 4f**). While all the major FeRB tend to increase and become enriched in plaque over time, the different patterns suggest that there are distinct FeRB niches.

Both FeOB and FeRB are also enriched in the rhizosphere (Fig. 3, Fig. 4). Although the timing and magnitude of enrichment differs somewhat from plaque, the enrichment suggests that these organisms, particularly the FeOB, could also be associated with iron oxyhydroxide formation in the rhizosphere (satellite plaque).

#### Correlations between FeOB, FeRB, and plaque iron

Both FeOB and FeRB were consistently enriched and abundant in the plaque. To better understand their relationships with iron geochemistry, we looked for correlations between predominant plaque organisms and plaque Fe (**Fig. 5, Fig. S3**). Plaque Fe content was positively correlated with the relative abundance of the top FeOB *Gallionella* OTU7 (rho=0.47, p=0.09, **Fig. 5c**) and top FeRB *Anaeromyxobacter* OTU0 (rho=0.53, p=0.05, **Fig. 5a**). Because the data are from 12 paddies that vary in geochemistry, we also plotted trends within individual paddies in **Fig. 5b and 5d**. In cases where data were available for multiple time points in specific paddies, the relative abundance of both *Gallionella* OTU7 and *Anaeromyxobacter* OTU0 corresponded to higher plaque Fe in each paddy.

**Figure 5.**
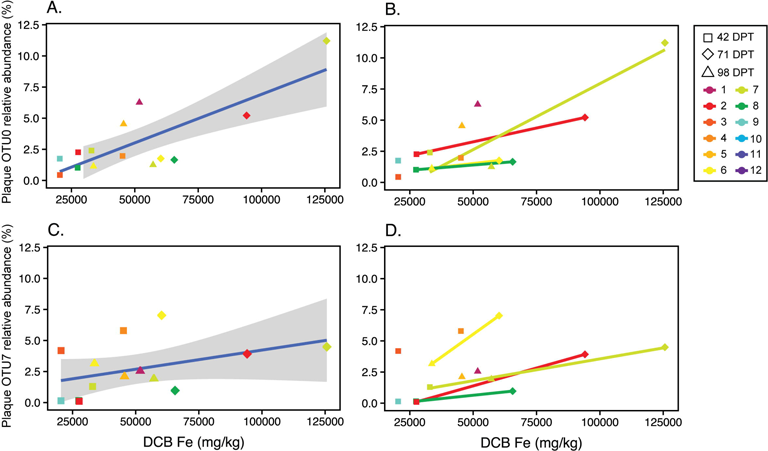
Comparison of selected FeRB and FeOB abundance with plaque iron content. Scatter plots show plaque relative abundance of *Anaeromyxobacter* OTU0 vs. plaque Fe (A, B), and of *Gallionella*/*Ferrigenium* OTU7 and plaque Fe (C, D). Points are colored by paddy and shapes represent sampling time (DPT). In A and C, all plaque data are shown and the blue line shows linear regression and gray shaded region shows 95% confidence interval. In B and D, only paddies with data from multiple time points are shown and temporal trends are plotted by paddy.

#### FeOB genomes reconstructed from rhizosphere and bulk soil metagenomes

To investigate the metabolic capabilities of FeOB, we sequenced two metagenomes from bulk soil (9BB) and rhizosphere (9BH), which were sampled from the same paddy in the same year at early reproduction (42 DPT). Based on our 16S rRNA gene analyses, these two samples include all major FeOB and FeRB OTUs. Indeed, both metagenomes yielded high quality genomes of FeOB and FeRB (>90% completeness, <5% contamination) that reflected taxa identified by 16S analysis (**Table 1)**. Here we briefly describe the FeRB and then focus on analyses of FeOB to assess their potential metabolic influences on biogeochemical cycling as well as plant interactions.

**Table 1.**
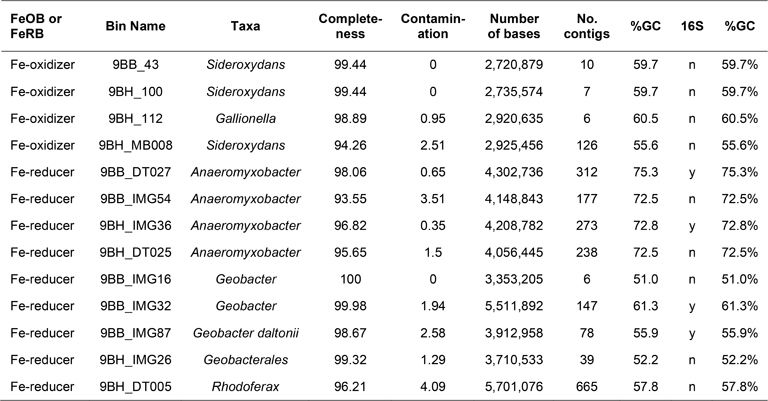
Metagenome-assembled genome bins.

| FeOB or FeRB | Bin Name | Taxa | Completeness | Contamination | Number of bases | No. contigs | %GC | 16S | %GC |
| --- | --- | --- | --- | --- | --- | --- | --- | --- | --- |
| Fe-oxidizer | 9BB_43 | *Sideroxydans* | 99.44 | 0 | 2,720,879 | 10 | 59.7 | n | 59.7% |
| Fe-oxidizer | 9BH_100 | *Sideroxydans* | 99.44 | 0 | 2,735,574 | 7 | 59.7 | n | 59.7% |
| Fe-oxidizer | 9BH_112 | *Gallionella* | 98.89 | 0.95 | 2,920,635 | 6 | 60.5 | n | 60.5% |
| Fe-oxidizer | 9BH_MB008 | *Sideroxydans* | 94.26 | 2.51 | 2,925,456 | 126 | 55.6 | n | 55.6% |
| Fe-reducer | 9BB_DT027 | *Anaeromyxobacter* | 98.06 | 0.65 | 4,302,736 | 312 | 75.3 | y | 75.3% |
| Fe-reducer | 9BB_IMG54 | *Anaeromyxobacter* | 93.55 | 3.51 | 4,148,843 | 177 | 72.5 | n | 72.5% |
| Fe-reducer | 9BH_IMG36 | *Anaeromyxobacter* | 96.82 | 0.35 | 4,208,782 | 273 | 72.8 | y | 72.8% |
| Fe-reducer | 9BH_DT025 | *Anaeromyxobacter* | 95.65 | 1.5 | 4,056,445 | 238 | 72.5 | n | 72.5% |
| Fe-reducer | 9BB_IMG16 | *Geobacter* | 100 | 0 | 3,353,205 | 6 | 51.0 | n | 51.0% |
| Fe-reducer | 9BB_IMG32 | *Geobacter* | 99.98 | 1.94 | 5,511,892 | 147 | 61.3 | y | 61.3% |
| Fe-reducer | 9BB_IMG87 | *Geobacter daltonii* | 98.67 | 2.58 | 3,912,958 | 78 | 55.9 | y | 55.9% |
| Fe-reducer | 9BH_IMG26 | *Geobacterales* | 99.32 | 1.29 | 3,710,533 | 39 | 52.2 | n | 52.2% |
| Fe-reducer | 9BH_DT005 | *Rhodoferrax* | 96.21 | 4.09 | 5,701,076 | 665 | 57.8 | n | 57.8% |

#### FeRB genomes

The metagenomes yielded nine high quality FeRB genomes, which were identified as *Anaeromyxobacter*, *Geobacter*/Geobacterales, and *Rhodoferax*. While most (6 of 9) were reconstructed from the bulk soil, one genome for each FeRB taxonomic group was recovered from the rhizosphere (**Table 1**). All of these FeRB genomes contained genes for oxygen respiration: *Anaeromyxobacter* and *Rhodoferax* genomes all encoded genes for both aa_3_- and cbb_3_-type cytochrome *c* oxidases while the *Geobacter* encoded either aa_3_ or cbb_3_ types. This suggests that all of the FeRB are facultative anaerobes.

*Rhodoferax* is known as an FeRB, but there is one recent report of Fe oxidation by *Rhodoferax* MIZ03 ((66); 81.7% ANI with *Rhodoferax* 9BH_DT005). The *Rhodoferax* genome from the rhizosphere includes genes that may catalyze either iron oxidation or reduction. These are homologs of the outer membrane-associated decaheme cytochrome MtoA/MtrA and porin MtoB/MtrB. In the FeRB *Shewanella oneidensis*, MtrA has also been shown to both reduce iron and take up electrons from an electrode (67). In the FeOB *Sideroxydans* ES-1, MtoA is an iron oxidase (68, 69). Phylogenetic analysis of the *Rhodoferax* 9BH_DT005 MtoA/MtrA homolog shows that it clusters with MtoA sequences from Gallionellaceae FeOB (**Fig. S4**). Altogether, this suggests that there is a possibility that the rice rhizosphere *Rhodoferax* may be a facultative FeRB or FeOB.

#### FeOB genomes

The metagenomes yielded four high quality FeOB genomes–three *Sideroxydans* and one *Gallionella*, classified based on a synthesis of GTDB results, average amino acid identity (AAI)/average nucleotide identity (ANI) (**Supp. Table S3**), and a phylogenetic tree of concatenated ribosomal protein sequences (**Fig. 6**). The *Sideroxydans* bins from the bulk soil (9BB_43) and rhizosphere (9BH_100) are 100% identical by ANI and cluster with *Sideroxydans lithotrophicus* ES-1 (81% ANI/82% AAI). The rhizosphere genome bin 9BH_112 is closely related to the rice paddy soil isolate *Ferrigenium kumadai* An22, (90.1% ANI/91.5% AAI); however, GTDB classifies AN22 and bin 9BH_112 as *Gallionella,* which is also confirmed by our phylogenetic analysis based on 13 concatenated ribosomal protein sequences (**Fig. 6**; details in methods and (35)). Thus, going forward, we refer to bin 9BH_112 as *Gallionella* 9BH_112 with the understanding that it is closely related to *F. kumadai* An22. The unclassified Gallionellaceae 9BH_MB008 is more distant from isolates: 79.5% ANI to *Sideroxydans* ES-1, 79.4% ANI to *Ferrigenium* An22, 79.0% ANI to *Ferriphaselus* R-1. GTDB classifies 9BH_MB008 as genus “Palsa-1006,” which has no cultured representatives, and to our knowledge, this clade has not yet been described from a rice paddy environment. Our whole genome phylogenetic analysis determined that 9BH_Bin008 and the GTDB “Palsa-1006” fall within *Sideroxydans*, so we refer to this bin as *Sideroxydans* 9BH_MB008.

**Figure 6.**
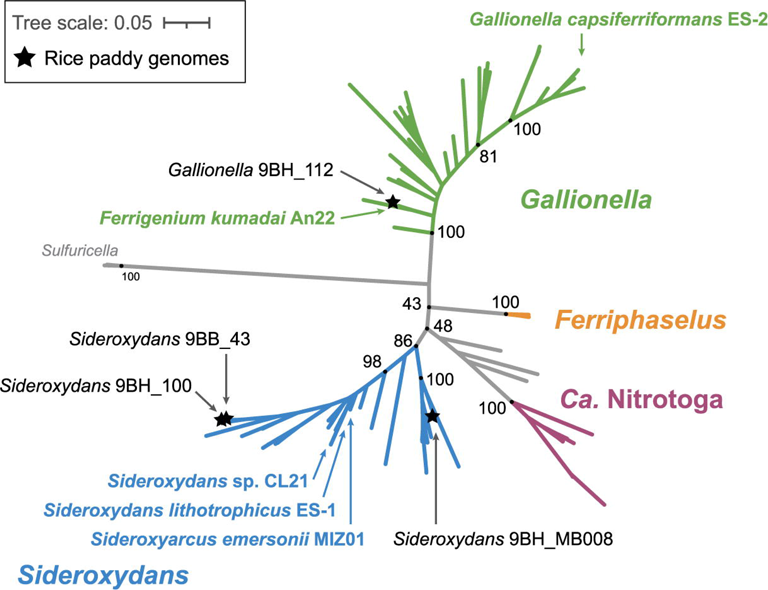
Maximum likelihood tree of Gallionellaceae FeOB genomes based on 13 concatenated ribosomal protein sequences. Genomes from this study are shown with a star, and isolate genomes are shown by arrows only. While *Gallionella* 9BH_112 is closely related to the rice paddy isolate *F. kumadai* AN22, the other genomes in this study are more distant from isolates (in *Sideroxydans*). Support values based on 1000 bootstraps. Further information on classification and tree construction, including all genomes in the tree, are available in Hoover et al. (29).

While none of the reconstructed Gallionellaceae genomes from 9BB or 9BH samples include 16S rRNA genes, we can use metagenomes from samples from the following year (2017) to help connect the genomes to 16S rRNA sequences. The *Sideroxydans* genomes 9BB_43 and 9BH_100 are practically identical (99.9% ANI, 100% AAI) with another genome reconstructed from the same sampling site a year later (9GH_6), which does have a 16S rRNA gene fragment. The 9GH_6 16S rRNA gene fragment shares 98.4% identity with a representative *Sideroxydans* OTU2 16S rRNA gene fragment. In all, the phylogenetic analyses suggest that genome reconstruction yielded genomes that are representative of the major FeOB detected by 16S analyses, *Sideroxydans* OTU2 and *Gallionella* OTU7, plus one additional *Sideroxydans* genome.

#### FeOB energy metabolisms - electron donors and acceptors

We analyzed Gallionellaceae bins to explore the FeOB for biogeochemical contributions, including Fe, C, N, P, and S cycling. Genes and metabolisms are summarized in **Figure 7A**. *Iron oxidation.* All four 9BB/9BH Gallionellaceae genomes have genes for microaerophilic iron oxidation. The genomes contain the iron oxidase gene *cyc2*, which is characteristic of most Gallionellaceae (35). They lack *mtoAB*, an outer membrane multiheme cytochrome-porin complex associated with iron oxidation in *S. lithotrophicus* ES-1 (68–70). Each genome has the potential for oxygen respiration, with at least one set of terminal oxidase genes, *ccoNOP, coxAB,* and/or *cydAB*.

**Figure 7.**
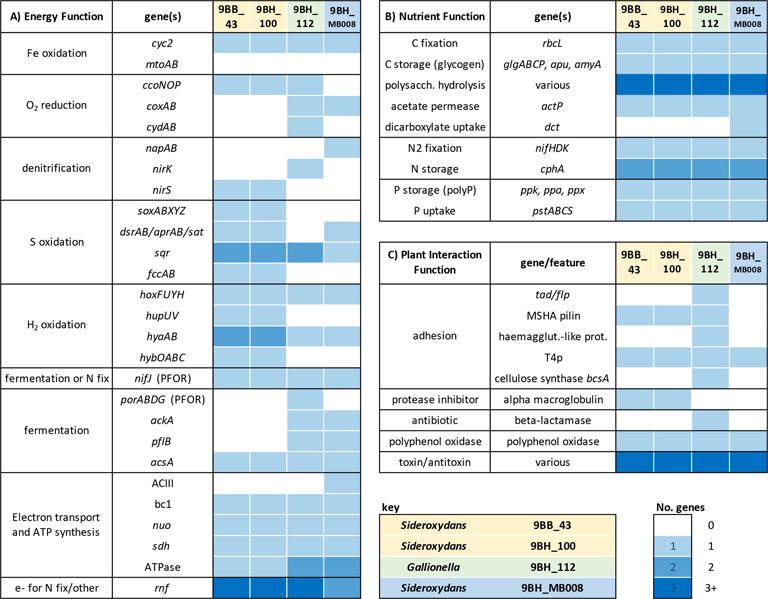
Summary of selected functional genes in Gallionellaceae FeOB genomes. Boxes are colored by number of genes (“No. genes”) or gene sets (if >1 gene mentioned) present in each genome. See Table S4 for polysaccharide hydrolysis genes. “N fix” = nitrogen fixation.

##### Denitrification

Only *Sideroxydans* 9BH_MB008 has the potential for dissimilatory nitrate reduction, with *napAB* in its genome. The *Sideroxydans* 9BB_43 and 9BH_100 genomes encode nitrite reductase *nirS* while *Gallionella* 9BH_112 has the *nirK* nitrite reductase. These denitrification genes are not sufficient for growth using nitrate-dependent iron oxidation metabolism, though *Sideroxydans* 9BH_MB008 may be able to use denitrification for redox balance when O_2_ is limiting, as posited for Zetaproteobacteria (71).

##### Sulfur oxidation

All bins had genes for sulfur oxidation. In particular, *Sideroxydans* 9BB_43 and 9BH_100 have genes for multiple sulfur oxidation pathways, including thiosulfate oxidation (*soxABXYZ)* and sulfide oxidation (*dsrAB/aprAB/sat*, *fccAB, sqr*). *S. lithotrophicus* ES-1 similarly has *sox* and *dsr* genes, and is able to grow by thiosulfate oxidation. Based on previous phylogenetic analysis, the DsrAB from ES-1 and the four 9BB/9BH Gallionellaceae genomes cluster with reverse DSR (rDSR) (35).

##### Hydrogen

All four Gallionellaceae bins have the potential for hydrogen oxidation using a variety of hydrogenase genes. All have *hoxFUYH* and *hyaAB,* while the *Sideroxydans* genomes 9BB_43 and 9BH_100 also have *hupUV* and *hybOABC.* Hydrogenase genes are common amongst FeOB (e.g. (35, 72, 73)) and other bacteria, and may play a role in providing reducing equivalents for carbon and nitrogen fixation since the Fe(II)/Fe(III) redox potential is relatively high and requires reverse electron transport to produce NADH and ferredoxin. The variety of hydrogenases implies a range of roles, which may also include growth on hydrogen as well as fermentation, with hydrogen formation serving as an electron sink (see below).

##### Electron transport chain

Genes for components of the rest of the electron transport chain are present including Complex III (bc1) NADH dehydrogenase, and ATPase(s). Genes for alternative complex III (ACIII) were not found in the *Sideroxydans* 9BB_43 and 9BH_100 or *Gallionella* genomes but were present in *Sideroxydans* 9BH_MB008.

##### Fermentation/”anaerobic” metabolism

The Gallionellaceae genomes have a number of genes associated with anaerobic metabolisms, possibly for fermentation. *Gallionella* 9BH_112 and *Sideroxydans* 9BH_MB008 both have pyruvate formate lyase, which converts pyruvate to formate and acetyl CoA. In addition, all four Gallionellaceae encode genes for pyruvate ferredoxin/flavodoxin oxidoreductase (PFOR), which would generate acetyl coA and reduced ferredoxin or flavodoxin. The four genomes also encode acetyl CoA synthetase which can generate ATP during the conversion of acetyl CoA to acetate. In some cases the PFOR genes are adjacent to *hox* hydrogenase genes, which may be used to evolve H_2_ for redox balance (e.g. (74)). As these Gallionellaceae have genes for glycogen formation and usage, this altogether suggests the ability to ferment glucose into acetate, which would allow energy generation for survival of anoxic conditions. It has also been shown that PFOR can be used in place of pyruvate dehydrogenase under oxic, but highly reducing conditions as cells shift toward using ferredoxin-dependent enzymes, rather than NADH, with the lower redox potential ferredoxin capturing more energy and reducing power (75).

##### Summary

Analysis of energy metabolism genes show that the Gallionellaceae genomes include genes for aerobic iron oxidation as well as sulfur oxidation, hydrogen oxidation, fermentation, and in some cases, denitrification. Thus, overall, the FeOB show metabolic flexibility that could serve to help them survive across chemical gradients or fluctuating redox conditions. The ability to use multiple substrates can provide additional energy and reducing power for biosynthetic reactions such as C and N fixation.

#### Nutrient (C, N, P) acquisition, fixation, storage, and usage

As the FeOB are closely associated with roots, they may help promote plant growth by providing nutrients. To evaluate this, we looked for genes involved in the fixation, storage, and release of nutrients (**Fig. 7B**). In addition, we took note of genes that suggest utilization of plant exudates and polysaccharides. As Gallionellaceae are typically considered autotrophic, such genes could further link these FeOB to the plant rhizosphere environment.

##### Carbon fixation and storage

All Gallionellaceae isolates can grow autotrophically (28, 32, 33, 76–78), and like these isolates, all four Gallionellaceae genomes in this study include the Rubisco gene *rbcL* and other genes for the complete Calvin Benson Bassham pathway for carbon fixation. The genomes also include genes for glycogen formation and degradation, which allow cells to store carbon and energy.

##### Organic carbon utilization

While Gallionellaceae isolates are not known for heterotrophy, the 9BB and 9BH Gallionellaceae genomes contain a number of genes for organic carbon utilization, including polysaccharide degradation, and proteinases, and organic uptake. We identified carbohydrate-degrading and production genes using the DRAM software-based classification of genes according to the Carbohydrate-Active enZYmes (CAZY) Database (**Table S4**; (79)). The four genomes contain numerous genes encoding enzymes for degradation of polysaccharides, including carbohydrate esterases and many glycoside hydrolases. We also noted numerous peptidases according to the MEROPS database, also using DRAM (https://www.ebi.ac.uk/merops/). Further work will be needed to confirm the target substrates for the genes identified above. *Sideroxydans* 9BH_MB008 encodes a dicarboxylate transporter, and all four genomes encode acetate permease. The use of organics is unusual in Gallionellaceae, as most isolates are autotrophic, except *S. lithotrophicus* CL21, an isolate from a peatland that can utilize lactate alongside Fe(II) (80). Altogether these results suggest that these Gallionellaceae are able to take advantage of active C cycling in the soil/rhizosphere and utilize plant- and microbial-derived organic carbon, and therefore grow as mixotrophic FeOB (as in ref (80, 81)).

##### Nitrogen fixation and storage

All four Gallionellaceae genomes have nitrogen fixation genes, including key genes *nifHDK,* as well as genes to produce the nitrogen storage compound cyanophycin and the Rnf complex, which provides electrons to nitrogenase by reducing ferredoxin via NADH (82, 83). These genes are not present in all Gallionellaceae genomes, likely because nitrogen fixation is an energy-intensive process that would be difficult to support using the limited energy available from chemolithoautotrophic iron oxidation. However, N_2_ fixation and storage would give FeOB a competitive advantage at the root surface, as the cells could help supply fixed nitrogen to the plant, which is N-limited due to increasing N demand with plant growth. Past work has shown that N fixation occurs in flooded paddies mainly at the heading stage in rice (84, 85), which coincides with when *Gallionella/Ferrigenium* OTU7 begins to increase in abundance in the plaque.

##### Phosphorous uptake and storage

Like all Gallionellaceae, the four 9BB and 9BH FeOB genomes have genes to take up inorganic phosphate. They also have genes for polyphosphate formation and utilization, allowing them to store phosphate. This storage would be useful as dissolved phosphate levels near the root may fluctuate as phosphate is adsorbed onto iron plaque. The four FeOB genomes also include a number of genes with annotations associated with organic phosphorous utilization, but a full pathway was not observed. Thus, there may be the potential for the FeOB to use organic phosphorous sources, but this is yet to be confirmed.

#### What other genes suggest adaptations to rice paddy soil and the root environment?

Because plaque is formed at the root surface, any plaque-forming microbes must occupy the rhizoplane and interact directly with plant roots. The Gallionellaceae genomes include various genes that could be involved in colonization and attachment to roots (**Fig. 7C**). The *Gallionella* 9BH_112 genome includes a number of adhesion genes that could mediate direct interactions with the plant root. *Gallionella* 9BH_112 has genes for the “widespread colonization island” that encodes for Tad/Flp pilin, which is responsible for tight attachment to surfaces. This includes flp-1, which encodes the structural component of the pilin as well as genes to secrete the pilin and assemble the pilus. These pili mediate surface adherence and biofilm formation, and have been shown to be important for colonization (86).

Another adhesive pilin, the mannose-sensitive hemagglutination (MSHA) pilin cluster, is encoded by *Sideroxydans* 9BB_43 and 9BH_100 and *Gallionella* 9BH_112. MSHA is involved in virulence in vibrio (87) and has also been shown to mediate non-virulent attachment of *Pseudoalteromonas tunicata* to the green alga *Ulva australis* (88). The *Gallionella* 9BH_112 genome also encodes a large (3638 aa) haemagglutinin-like protein adjacent to a gene encoding a hemolysin activation/secretion protein. The homologs in *Bordetella pertussis* encode FhaC, an outer membrane transporter, which secretes FhaB, a filamentous haemagglutinin that mediates adhesion to host tissues (89).

The various adhesion mechanisms would enable attachment to roots. After this initial attachment, the FeOB cells would form biofilms. The genomes include a variety of genes to help build biofilms, including polysaccharide synthesis. Notably, *Gallionella* 9BH_112 encodes cellulose synthase, and cellulose fibrils are known to aid in *Rhizobium* and *Agrobacterium* cell attachment to roots (90). *Gallionella* 9BH_112 and two *Sideroxydans* genomes 9BB_43 and 9BH_100 also encode a number of genes associated with capsular polysaccharide synthesis.

Colonization also requires the microbes to overcome plant immunity defenses. The *Sideroxydans* genomes 9BH_43 and 9BH_100 both include a gene encoding alpha 2 macroglobulin, a protease inhibitor that is a factor in virulence and colonization (91). The alpha 2 macroglobulin gene is adjacent to a penicillin-binding protein 1c, similar to *Salmonella enterica* ser. typhimurium and other bacteria. These two are thought to work together, with alpha 2 macroglobulin inhibiting proteases when cell walls are ruptured while the penicillin-binding protein repairs cell wall peptidoglycan (91). Plants produce polyphenols, which are toxic to bacteria (92); all four genomes encode polyphenol oxidases which help defend against these antimicrobial compounds.

Other genes that may be involved in interactions with plants and other rhizosphere microbes include genes for antibiotics, and toxin/antitoxins. In all, the Gallionellaceae FeOB appear to be well-equipped to colonize plant roots, deploy immunity or antibiotic defenses as needed, contribute nutrients to plants, and utilize plant exudates in their own nutrition.

### Conclusions and Implications

Our overall goal was to determine whether iron-oxidizing bacteria are associated with iron plaque and gain insight into their metabolism, dynamics, and potential plant interactions, toward understanding FeOB roles in plaque formation. In contrast to iron reduction, microbial iron oxidation is often an invisible process in the environment because abiotic processes can also oxidize Fe(II) and it can be difficult to distinguish biotic and abiotic effects. However, it is important to do so because microbial iron oxidation also drives C, N and P cycling and potentially plant processes. Here we reveal the dynamics of FeOB and FeRB in rice root plaque and rhizosphere across an entire growing season. We show that Gallionellaceae, which are well-known chemolithotrophic FeOB, are specifically enriched in both plaque and rhizosphere microbial communities and have the genetic potential to aerobically oxidize Fe(II), cycle nutrients, colonize roots, and contribute to plant nutrition.

We detected at least two different Gallionellaceae; 16S rRNA gene analyses revealed that *Sideroxydans* dominated early in the season while a *Gallionella* related to *Ferrigenium* increased over time, as plaque increased. Genomics revealed an additional, phylogenetically distinct *Sideroxydans* (9BH_MB008) that was not distinguished during 16S analyses. Both the temporal dynamics and the coexistence of the different Gallionellaceae suggest distinct niches. *Sideroxydans* are facultative iron oxidizers, shown to grow on other substrates such as thiosulfate (69, 93) and H_2_ (80). At the same time, there are signs that *Sideroxydans* are primarily FeOB, as *S. lithotrophicus* ES-1 highly expresses iron oxidation genes during thiosulfate oxidation (69). The *Sideroxydans* may use their alternate (non-Fe) energy metabolisms to thrive and create biomass in low Fe conditions, putting them in position to quickly colonize rice roots early in the season.

Gallionellaceae are known as chemolithoautotrophs, but they are commonly found in soils and sediments, and have been found to thrive in organic-rich environments like wetlands/fens (33, 94–96). Accordingly, Gallionellaceae, including the rice paddy genomes in this study, have genes for organic uptake and utilization, i.e. heterotrophy. This suggests that Gallionellaceae in fact can grow mixotrophically using both Fe(II) and organics, taking advantage of plant exudates as a carbon source. The rice paddy Gallionellaceae have genes that further suggest a specific niche at the root surface, as they have genes for plant interactions, including colonization, immunity, uptake/utilization of plant organics, and nutrient fixation. *Gallionella* 9BH_112 is especially rich in these genes, which may explain why it increases in relative abundance as the plant matures and root mass increases. Plants are more likely to tolerate root-associated microbes that help promote plant growth, and indeed, the FeOB could provide nutrition to the rice plants. All of the Gallionellaceae genomes encode nitrogen fixation, which is otherwise a relatively unusual trait in the Gallionellaceae (< 20% of other Gallionellaceae genomes; (35)). As N fixation is an energy-intensive process, this further supports the specificity to the paddy soil environment, as Gallionellaceae have the capacity to contribute to plant nutrition and growth. Our work shows that the rhizoplane/rhizosphere hosts these microaerophilic FeOB, whose activity can produce iron oxyhydroxides. The increasing relative abundance corresponds to increasing oxyhydroxides, and thus in total, the evidence suggests these Gallionellaceae FeOB are specifically suited to the rhizosphere/rhizoplane niche and very likely contribute to plaque formation.

Alongside the FeOB, we also find FeRB enriched in the plaque, and both FeOB and FeRB increase in relative abundance over time. While it may be surprising that FeRB would occupy the oxic zone and increase over time as oxygen levels increase, the FeRB appear to be aerotolerant, and it makes sense that FeRB would increase with increasing iron oxyhydroxide. This suggests an active iron cycle within the plaque, as well as the rhizosphere and bulk soil, where both FeOB and FeRB are also found. This co-existence of FeOB and FeRB has also been observed in other iron-rich rhizospheres of other wetland plants (7, 97). The production of iron plaque is likely due to the concerted action of both oxidizers and reducers, as follows. In the more reducing bulk soil, iron reduction coupled to oxidation of organics fuels the formation of dissolved Fe^2+^. Mass flow toward the roots pulls the Fe^2+^ into the more oxic rhizosphere and root surface, where FeOB catalyze Fe^2+^ oxidation to Fe(III) oxyhydroxides. The coexistence of FeOB and FeRB suggest further redox cycling, and continued oxidation and reduction that results in dynamic remodeling of rhizosphere and rhizoplane oxyhydroxides (i.e. plaque). Thus, the formation of plaque requires *both* reducing and oxidizing conditions, and *both* FeOB and FeRB. This dynamic redox conceptual model has implications for oxyhydroxide-associated elements, and therefore plant interactions with elements like P and As. While oxyhydroxides may collect and temporarily sequester phosphate and arsenate, iron reduction in the plaque will release these elements, with consequent impacts on plant health and food safety. Further temporal studies will be needed to determine if iron-cycling dynamics within rice root plaque ultimately cause greater uptake of phosphorous and metals into plant and grain. However, this work suggests the rapid iron redox cycling between FeOB and FeRB could contribute to the plaque behaving as a dynamic source of nutrients and contaminants.

## Methods

### Rice paddy experiment

Samples for microbial and geochemical analyses were collected in 2016, described by (42) and (43), and in 2017 from the University of Delaware Rice Information, Communication, and Education (RICE) Facility. The RICE facility is an outdoor facility on the UD Farm in Newark that consists of 30 rice paddy plots (2×2 m) in which each soil-filled paddy accommodates 49 plants (7 rows of 7) and is outfitted with a bilge pump connected to a float switch and an irrigation line to control water level (Figure S1). In brief, rice (*Oryza sativa* L. ‘Jefferson’) for this experiment was grown each year in 12 of the paddy mesocosms that contained non-amended native soil or native soil amended with Si at identical Si loading rates of 5 Mg Si/ha with either rice husk, charred rice husk, or calcium silicate/silicic acid as the Si source (3 replicate paddy mesocosms per treatment). Paddies were amended prior to planting in 2015 and did not receive further Si amendments in 2016 (44) or in 2017. Treatments had less of an impact than redox status and microhabitats on the microbial community (98, 99); therefore, the present study focused on diversity and abundance of microbes associated with Fe cycling over the rice life cycle and data are averaged across all four treatments. Prior to planting in 2016 and in 2017, only roots from the previous year were tilled into the soil and paddies were fertilized with 112 kg N/ha as urea and 135 kg K_2_O/ha as KCl based on soil testing and fertility recommendations. The silty clay loam soil contained 1.9 (±0.5) g/kg acid ammonium oxalate extractable Fe and 12.0 (±1.8) g/kg dithionite-citrate-bicarbonate extractable Fe as measures of amorphous and crystalline Fe (average ± standard deviation, n=12) (100). The soil pH was 6.1 (±0.5) and the soil organic matter was 2.4 (±0.6) % as measured by loss on ignition. During growth, paddies were kept flooded at transplanting until grain maturity when paddies were drained before harvest. In 2016, paddies were drained at 96 days post transplant (DPT) and in 2017, paddies were drained at 100 DPT.

### Porewater chemical analyses

Weekly porewater data for 2016 are described in detail in a separate manuscript (43). In brief, porewater samples were collected weekly from June 8^th^, 2016 to September 6^th^, 2016 with Rhizon samplers (polyethersulfone, 0.15 μm pores, Soilmoisture Equipment Corp., Goleta, CA). Porewater samples were aliquoted for colorimetric measurement of dissolved Fe(II) by the ferrozine method (101), and separate porewater samples were acidified with 2% HNO_3_ prior to analysis for total Fe by ICP-OES (Thermo Iris Intrepid II XSP Duo View ICP). Because porewater sampling did not occur on the same days as microbial sampling, porewater values used for comparison are an average of the closest porewater values preceding (by no more than 7 days) and following (by no more than 7 days) the microbial sampling event, except in the case of the harvest time point (98 DPT) in which only the preceding porewater time point (96 DPT) was used. Porewater sampling dates averaged for comparison to microbial sampling events are summarized in **Table S5**.

### Root and soil sampling

During transplanting in May 2016 and 2017, five seedlings for each paddy mesocosm (60 plants total per year) were individually placed into 100 µm pore-size nylon mesh bags filled with paddy soil, according to (102) to define the rice rhizosphere. Different bag sizes were used to accommodate plant growth and were (diameter x depth) 10.2 cm x 20.3 cm for samples harvested at vegetative growth (20 days past-transplant (DPT)), 12.7 cm x 20.3 cm for early reproduction (42 DPT), and 15.2 cm x 31.8 cm for heading (71 DPT), grain ripening (88 DPT), and harvest (98 DPT). The five plants in mesh bags were randomly placed among 42 other transplanted seedlings.

To collect samples, bags were pulled from each paddy mesocosm at designated growth stages, and bulk soil was collected from the area surrounding the bag. Samples were immediately brought to the laboratory and separated into biospheres (plaque and rhizosphere) on ethanol-sterilized benchtops. Using sterilized instruments, rice roots were gently separated from the shoots. Roots were submerged in 25 mL 18 MΩ-cm sterile water (2016) or RNA later (2017) and vortexed twice to collect the rhizosphere. Bulk soil and rhizosphere samples were frozen at −20 °C until DNA extraction. Cleaned, rhizosphere-free roots were then divided into subsets, with half frozen prior to collecting the plaque, and half dried prior to DCB extraction of plaque Fe (described below). To collect the plaque for DNA extraction, thawed roots were transferred into 10 mL phosphate buffer solution and sonicated for 30 s twice (modified from (62)). Samples were centrifuged at 670 x g for five minutes to collect plaque, supernatant was decanted, and DNA was immediately extracted.

### Plaque iron and mineralogical analysis

Fe plaque was characterized to examine changes in the quantity of plaque Fe over the rice life cycle and Fe mineral composition at harvest according to established protocols (9, 103). For this, roots in collected bags (described above) were subject to dithionite-citrate-bicarbonate extraction (103), and total Fe was measured via ICP-OES. At harvest, an additional subset of soil-free roots (three root masses sampled diagonally across each paddy and pooled) were sonicated in water and filtered onto 0.2 µm nitrocellulose membranes for bulk analysis of plaque Fe mineral composition via Fe X-ray absorption fine structure (EXAFS) spectroscopy according to (9). Samples were analyzed at the Stanford Synchrotron Radiation Lightsource (SSRL) on beamline 11-2. There, incident energy was calibrated by assigning the first Fe inflection point from a standard Fe foil of 7112 eV, and two Fe K-edge EXAFS spectra were recorded for each sample in fluorescence with a Lytle detector. Spectra were averaged, normalized, background subtracted and fit by linear combination in Athena using reference spectra of known minerals found in Fe plaque including ferrihydrite, goethite, lepidocrocite, and siderite. Further details regarding geochemical analysis are available in (43).

### DNA extraction

For all bulk, rhizosphere, and plaque samples, DNA was extracted with the following modifications to the manufacturer’s instructions for the DNeasy PowerSoil DNA extraction kit (Qiagen). Powerbead solution (200 µL) was removed from bead-beating tubes and replaced with 200 µL phenol:chloroform:isoamyl alcohol 25:24:1 (v/v/v). The protocol was followed to manufacturer’s specifications for the PowerSoil kit until conditions were adjusted for column binding. At this step, equal parts lysate, solution C4, and 100% ethanol were homogenized and loaded onto the DNA binding column. After DNA binding, the column was washed with 650 µL 100% ethanol, followed by 500 µL solution C5. The column was centrifuged and dried, and DNA was eluted with molecular biology grade water (protocol based on personal communication with a Qiagen representative).

### 16S rRNA gene sequencing and analysis

Samples were submitted to the Joint Genome Institute for paired-end (2 × 300) Illumina MiSeq iTag sequencing of the 16S rRNA gene V4-V5 region, using primers 515F-Y and 926R (104). Raw sequences were de-multiplexed, quality filtered and clustered to 97% similarity with usearch, checking for chimeras, as part of the JGI pipeline. Feature tables and sequences were imported into QIIME 2 (105), and additionally filtered to exclude any features with a frequency of less than 10 sequences observed in the dataset that did not appear in at least two samples. Taxonomy was assigned to features using the qiime2 sklearn naïve bayes feature classifier, originally trained with SILVA database version 132 (https://www.arb-silva.de/fileadmin/silva_databases/LICENSE.txt). An initial report of the 16S rRNA data was given in reference (99). Data were imported into R (v3.6.2, (106)) and prepared for analysis using phyloseq (v1.28.0, (107)) with qiime2R (v9.99.13, (108)). Differential abundance analysis was assessed between plaque and bulk samples at individual timepoints using DESeq2 (109). Figures were built using phyloseq, the phyloseq wrapper for vegan (v2.5-6, (110)) and ggplot2 (v3.3.2, (111)).

### Metagenome sample selection, sequencing, and analysis

For metagenome sequencing, we selected samples that had sufficient DNA and good representation of major iron cycling OTUs. The majority of the data presented here are from a paired rhizosphere and bulk soil sample set from 2016, 9BH and 9BB, sampled at 42 DPT from Paddy 9, a “control” paddy that received no silica amendments. Metagenome sequencing was also performed on two rhizosphere samples from 2017, 9GH and 9IH, from Paddy 9 at 45 and 79 DPT. Here, the two 2017 samples are only used for linking 16S rRNA gene to the *Sideroxydans* genomes, but are available for further analyses on IMG. For the 2017 samples, all field and DNA extraction methods were as above, except that the rhizosphere sample was collected by vortexing roots in 25 mL of RNA later.

Metagenome sequencing was performed at the Joint Genome Institute (JGI) using an Illumina NovaSeq S4 (270 bp fragment, paired end 2×151). Initial data processing was done by JGI, including read correction by bbcms v 38.86, (112), assembly by metaSPAdes v. 3.14.1 (113), and annotation by IMG Annotation Pipeline v.5.0.20. Initial binning was performed by MetaBAT v2.15, evaluated using CheckM v1.1.3 (114). Medium and high quality genomes (>=50/90% completeness, <10/5% contamination; defined by the Genome Standards Consortium; (115)) were uploaded to IMG.

We performed additional binning on the KBase platform using MaxBin v2.2.4 (kb_maxbin v1.1.1) (116), CONCOCT v.1.1 (kb_concoct v1.3.4) (117), MetaBAT2 v1.7 (kb metabat v2.3.0) (118, 119), and then compared and optimized these bins using DASTool v1.1.2 (kb_das_tool v1.0.7) (120). Resulting genome bins were evaluated using CheckM v.1.0.18 (kb_Msuite v1.4.0) (114) and classified by GTDBtk v. 1.6.0 release 202 (kb_gtdbtk v1.0.0) (121, 122). We selected the highest quality genome of each known FeOB and FeRB taxa (>90% completeness, <5% contamination) from each sample (9BB and 9BH) for further analysis (Table 1). Genome similarity was analyzed by computing average nucleotide identity (ANI) by FastANI (123) in Kbase and average amino acid identity (AAI) by CompareM (124). We also assessed the phylogeny of Gallionellaceae FeOB by creating a concatenated gene tree. Thirteen ribosomal proteins (L19, L20, L28, L17, L9_C, S16, L21p, L27, L35p, S11, S20p, S6, S9) were aligned, trimmed, masked to remove regions with >70% gaps, and concatenated in Geneious v.10.2.6 (125). A maximum likelihood tree was constructed in RAxML-NG v1.0.3 (126) with the LG+G model and 500 bootstraps. Visualization and annotation of the final tree was done in iTOL (127). Final Gallionellaceae classifications, as described above, used the concatenated gene tree-based phylogeny and the scheme determined by Hoover et al., biorXiv/in review.

FeOB and FeRB bins selected for further analysis were annotated by RAST, DRAM (kb_DRAM v.0.1.2) (128), and FeGenie (129). For certain genes of interest, we further investigated close homologs by BLAST (130) against Uniprot database (131), by Blastp against functionally characterized sequences, and evaluation of alignments for conserved functional residues. The FeOB Cyc2, MtoA, and MtoB sequences were compared against *Sideroxydans lithotrophicus* ES-1 sequences Slit_0263, Slit_2497, and Slit_2496. For improved confidence in functional assignments, genomic context and synteny was evaluated in annotations, e.g. in RAST output.

To evaluate the potential function of the *Rhodoferax* Mto/Mtr homolog, we added the sequence to a tree of Mto and Mtr sequences from Hoover, et al. (35). The tree includes reference sequences of functionally verified MtoA, MtrA, and PioA along with additional Mto/Mtr sequences outlined by Hoover, et al. and Baker et al. (35, 132). Sequences were aligned, trimmed, and masked to remove gaps >70% in Geneious v.10.2.6 (125). A maximum likelihood tree was constructed using RAxML-NG v1.0.3 (126) with 500 bootstraps. The final tree was visualized and annotated using Iroki (133).

## Data availability

16S rRNA gene sequences are deposited at JGI DOI 10.25585/1488298 (iTagger processed and raw sequences), and NCBI under accession <u>PRJNA690162</u> (raw sequences). Metagenome sequences are available under NCBI Bioprojects PRJNA710504, PRJNA949703, PRJNA949704 and PRJNA1014123 and at IMG (genome IDs 3300042876, 3300044673, 3300044712, 3300045051; same DOI as above), including metagenome bins.

## Acknowledgements

We thank the Joint Genome Institute for funding and performing sequencing. We thank our field sampling team: Patrick Wise, Kristy Northrup, Weida Wu, Ayofela Dare, Kendall McCoach, Fred Teasley, Douglas Amaral, Ruifang Hu, Alesia Hunter, Julia O’Brien, and Heather Eby. We thank Olushola Awoyemi for help with genome analyses.

This work was supported by the National Science Foundation Grant Nos. 1350580 and 1833525, USDA NIFA Grant No. 2016-67013-24846, the DENIN Environmental Fellows program, the University of Delaware Doctoral and Dissertation Fellowships, and the Preston C. Townsend Biotechnology Fellowship. Sequencing for this project was completed by the Joint Genome Institute for project ID 503349. Use of the Stanford Synchrotron Radiation Lightsource, SLAC National Accelerator Laboratory, is supported by the U.S. Department of Energy, Office of Science, Office of Basic Energy Sciences under Contract No. DE-AC02-76SF00515.

This work is not a product of the United States Government or the United States Environmental Protection Agency. The author/editor is not doing this work in any governmental capacity. The views expressed are his/her own and do not necessarily represent those of the United States or the US EPA.

## Notes

### Competing Interest Statement

The authors have declared no competing interest.

### Summary of Updates

The manuscript has been revised throughout according to reviewer comments. Supplemental Figure S1 added to show rice paddy experiment.

