## Supplemental Figures for "Gallionellaceae in rice root plaque: metabolic roles in iron oxidation, nutrient cycling, and plant interactions"

Number of pages: 6

Number of figures: 4

Number of tables: 5

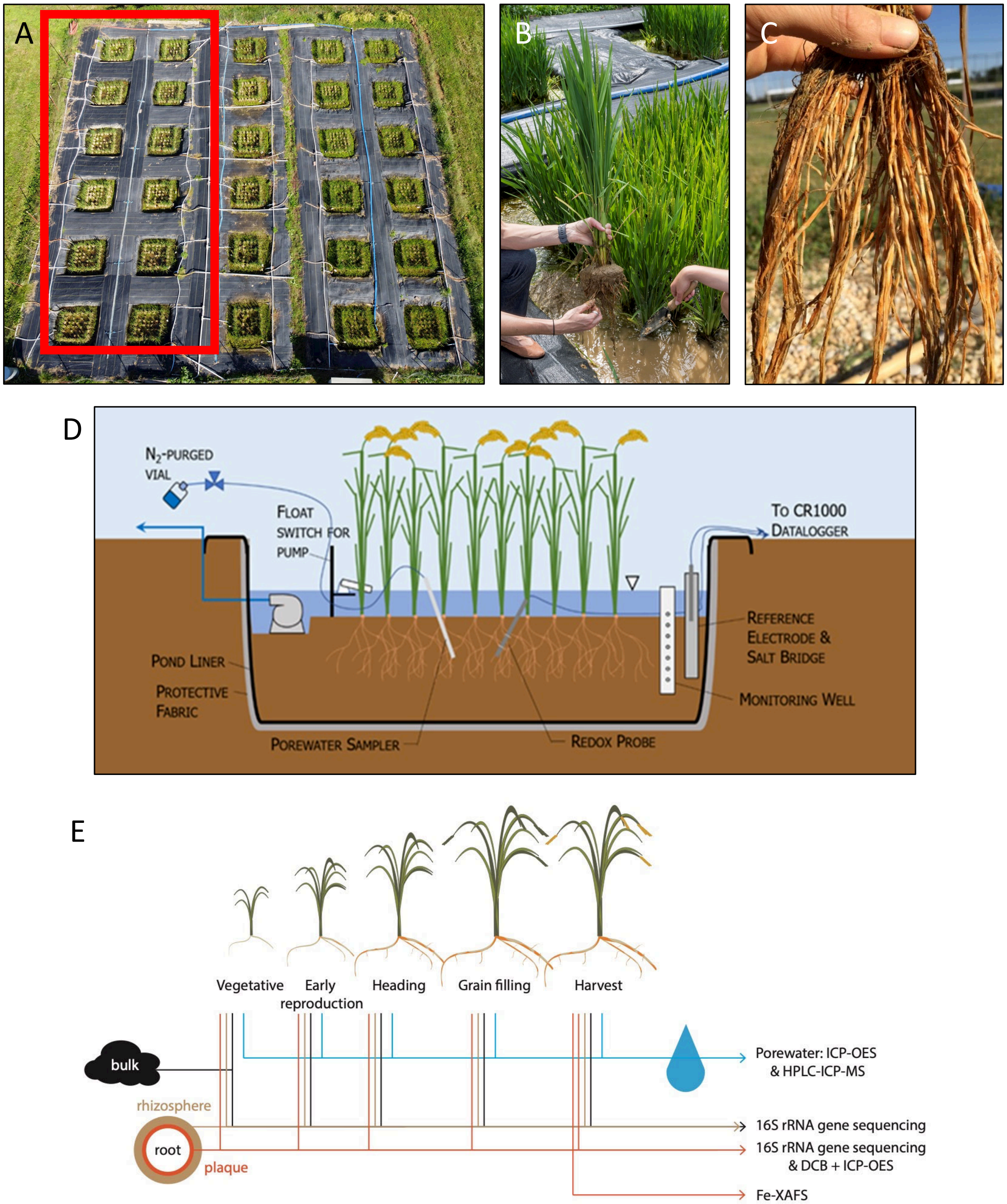


**Figure S1.** A) Overhead view of RICE facility, with the 12 paddies used in this study highlighted by the red rectangle. B) Individual rice plants were removed from each rice paddy throughout the growing season. C) Bulk soil and rhizosphere soil were removed from the plant to reveal the iron plaque. D) Schematic of an individual rice paddy. E) Timeline of plant and soil sampling, with specific analyses noted.

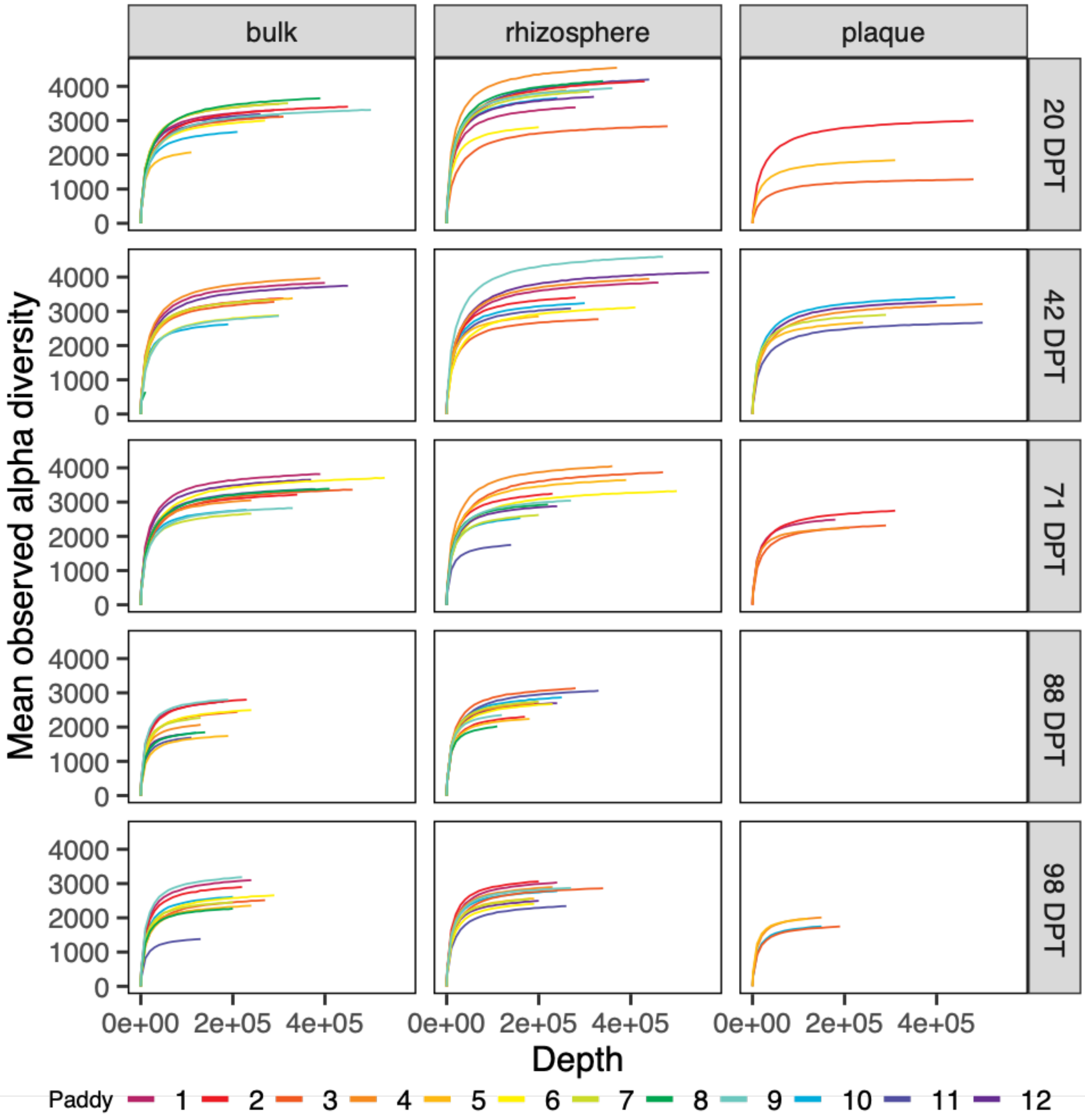


**Figure S2.** Observed alpha diversity rarefaction curves for each timepoint and biosphere. Rarefaction curves are colored by paddy.

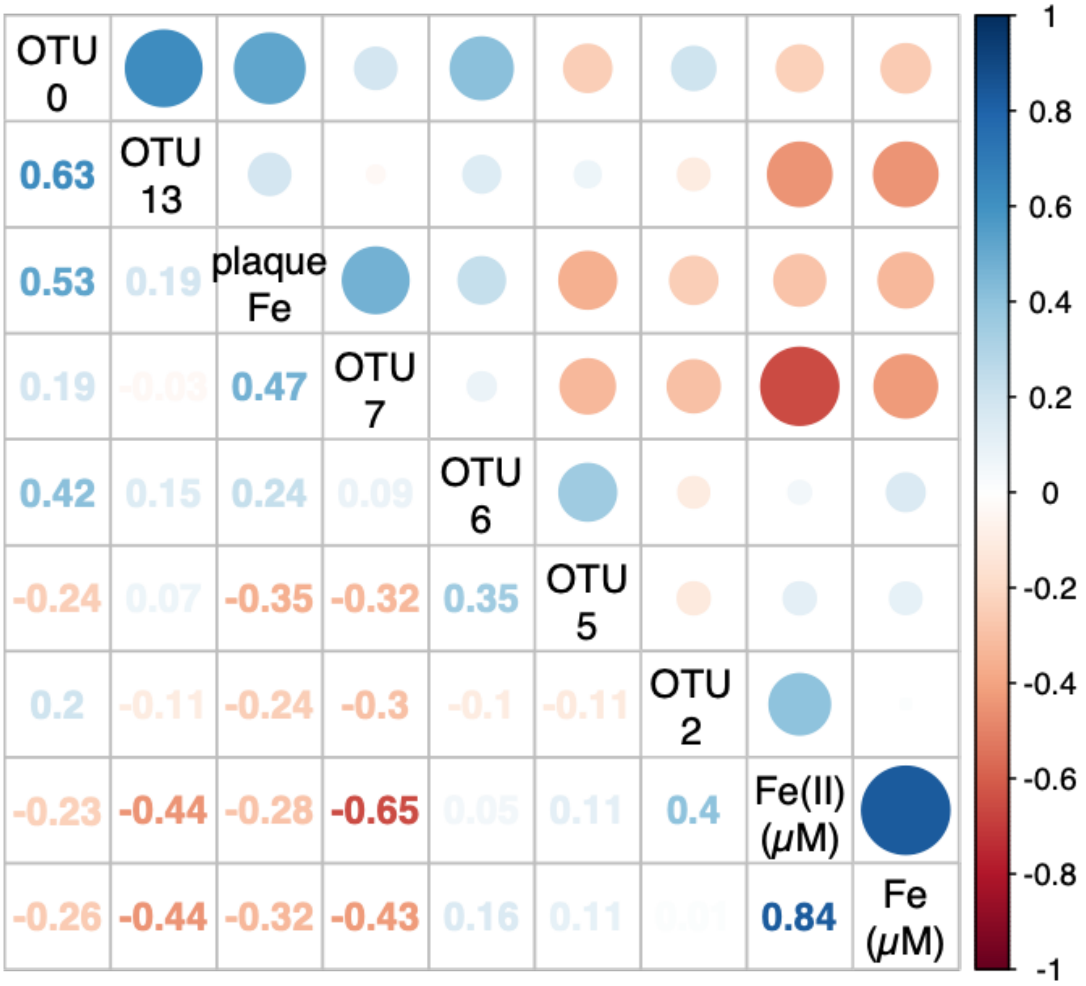


**Figure S3.** Spearman's rank correlation matrix of top Fe-cycling OTUs. Bubbles are sized and colored according strength and direction of Spearman correlation (see color scale), and rho is displayed on the lower left.

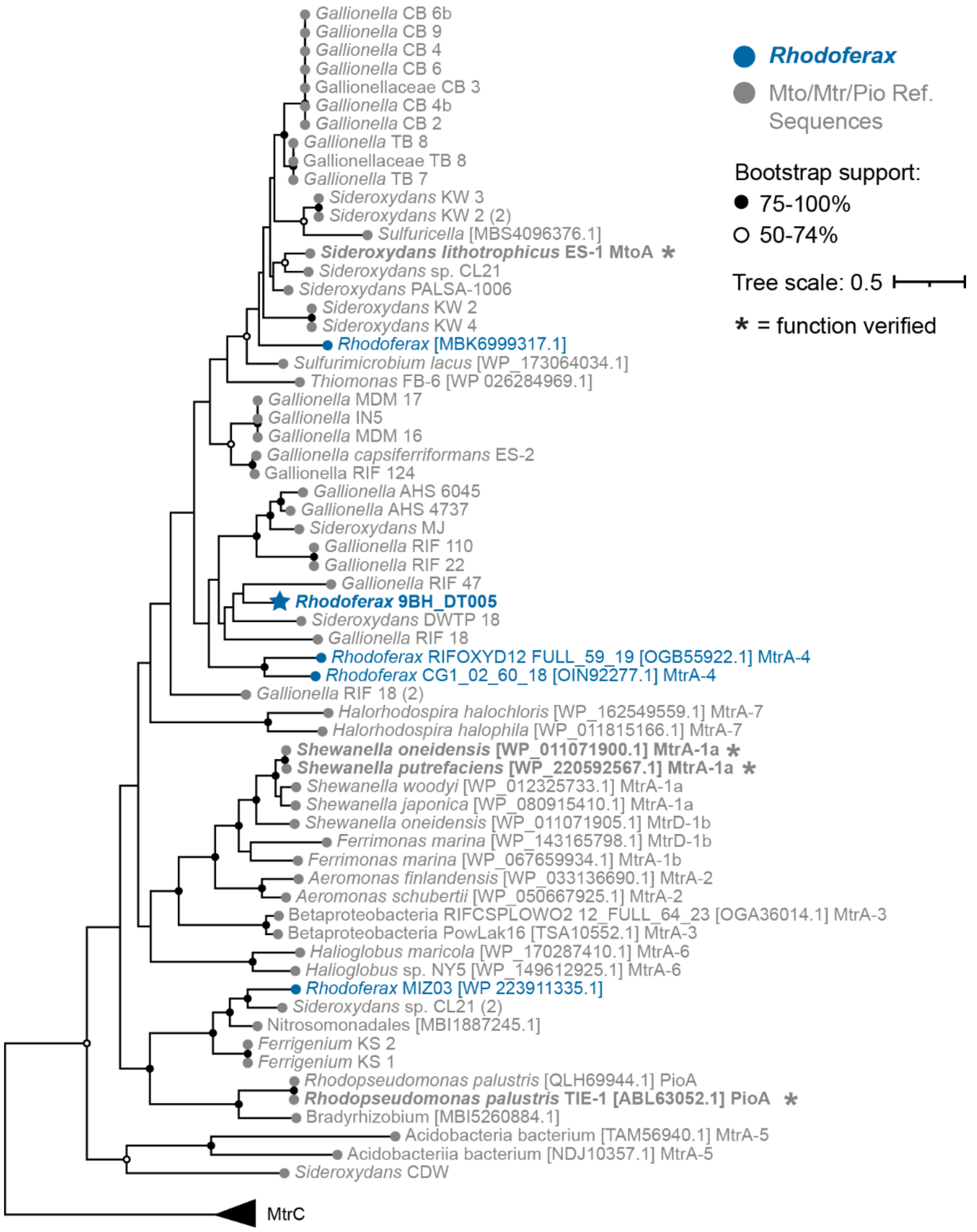


**Figure S4.** Maximum likelihood tree of MtoA/MtrA sequences showing that the *Rhodoferax* 9BH_DT005 sequence is closely related to functionally characterized iron oxidase MtoA sequences in *S. lithotrophicus* and within a clade of iron-oxidizing Gallionellaceae sequences. Support values based on 500 bootstraps. MtrA 1a, 1b, 2, 3, 4, 5, 6, and 7 indicate reference sequences from the seven MtrA groups defined by Baker, et al. (1).

### Supplemental Tables

Available as a separate Excel file.

**Table S1** OTU relative abundance and taxonomy

**Table S2** Top plaque OTUs

**Table S3** Genome average nucleotide and amino acid identity (ANI/AAI)

**Table S4** CAZY genes

**Table S5** Sampling time points for porewater and microbes
